# The interplay between sulfur assimilation and biodesulfurization phenotype in *Rhodococcus qingshengii* IGTS8: Insights into a regulatory role of the reverse transsulfuration pathway

**DOI:** 10.1101/2022.06.02.494632

**Authors:** Olga Martzoukou, Panayiotis Glekas, Margaritis Avgeris, Diomi Mamma, Andreas Scorilas, Dimitris Kekos, Sotiris Amillis, Dimitris G. Hatzinikolaou

**Affiliations:** Enzyme and Microbial Biotechnology Unit, Department of Biology, National and Kapodistrian University of Athens, Athens, Greece; Sector of Biochemistry and Molecular Biology, Department of Biology, National and Kapodistrian University of Athens, Athens, Greece; Laboratory of Clinical Biochemistry - Molecular Diagnostics, Second Department of Pediatrics, School of Medicine, National and Kapodistrian University of Athens, “P. & A. Kyriakou” Children’s Hospital, Athens, Greece; Biotechnology Laboratory, Sector of Synthesis and Development of Industrial Processes (IV), School of Chemical Engineering, National Technical University of Athens, Athens, Greece

**Keywords:** *Rhodococcus qingshengii* IGTS8, Biodesulfurization, Biocatalysis, Genetic engineering, Reverse transsulfuration, Sulfur metabolism

## Abstract

Biodesulfurization (BDS) is a process that selectively removes sulfur from dibenzothiophene and its derivatives. Several mesophilic natural biocatalysts have been isolated, harboring the highly conserved desulfurization operon *dszABC*. Even though the desulfurization phenotype is known to be significantly repressed by methionine, cysteine, and inorganic sulfate, the available information on the metabolic regulation of gene expression is still limited. In this study, scarless knockouts of the sulfur metabolism-related *cbs* and *metB* genes are constructed in the desulfurizing strain *Rhodococcus* sp. IGTS8. We provide sequence analyses for both enzymes of the reverse transsulfuration pathway and report their involvement in the sulfate- and methionine-dependent repression of the biodesulfurization phenotype, based on desulfurization assays in the presence of different sulfur sources. Additionally, the positive effect of *cbs* and *metB* gene deletions on *dsz* gene expression in the presence of both sulfate and methionine, but not cysteine, is uncovered and highlighted.

## Introduction

Microbial elimination of dibenzothiophene (DBT) and related organosulfur compounds, could allow for the biodesulfurization (BDS) of oil products by selectively removing sulfur from carbon-sulfur bonds, thus maintaining the calorific value of the fuel (1, 2). The process is mediated by the well-characterized mesophilic 4S metabolic pathway that is found in several genera, with the most prominent that of *Rhodococci* (3). The three BDS genes are organized in a plasmid-borne operon, *dszABC*, and encode for a DBT-sulfone monooxygenase (*dszA*), a 2-hydroxybiphenyl-2-sulfinate (HBPS) desulfinase (*dszB*), and a DBT monooxygenase (*dszC*), respectively. A fourth chromosomal gene, designated *dszD*, encodes for an NADH-FMN reductase that energetically supports the pathway. One of the major disadvantages in exploiting the biotechnological potential of the BDS process is the sulfate, methionine, and cysteine-mediated transcriptional repression of *dsz* genes through a putative repressor-binding site in the *P*_dsz_ promoter. The operon is de-repressed in the presence of organosulfur compounds such as DBT and dimethyl sulfoxide (DMSO), and Dsz enzymes are considered sulfate-starvation-induced (SSI) proteins (4–6). Mechanistic insight into the molecular regulation of *dsz* operon was gained recently with the identification of the TetR family activator DszGR, and the WhiB1 repressor, both derived from the desulfurizing bacterium *Gordonia* sp. IITR100 (6–8). Binding of DszGR to the promoter DNA induces an initial bend in *Gordonia* sp. *P_dsz_* region, but requires the integration host factor, IHF, which in turn plays a major role in promoter activity (9). Despite the high homology between *dszABC* operons of *G. alkanivorans* RIPI90A and *Rhodococcus* strain IGTS8, promoter sequences are only partially conserved (10). Moreover, a DszR activator also facilitated by IHF, has been reported for the activation of a σ^N-^dependent *dsz* promoter by metagenomic functional analysis (11). However, information on the global regulation of sulfur metabolism is still limited, and the sulfur assimilation pathways of *Rhodococci* had only been investigated *in silico* (12, 13). An exception is a very recent report that conducted comparative proteomics and untargeted metabolomics analyses in *Rhodococcus qingshengii* IGTS8 and proposed a working model for assimilatory sulfur metabolism reprogramming in the presence of DBT (4). Moreover, the effects of carbon and sulfur source on biotransformation of 6:2 fluorotelomer sulfonic acid were examined in *R. jostii* RHA1 (14). General aspects of carbon and nitrogen metabolism of oleaginous *Rhodococcus* spp. have been elucidated, due to the ability of this Rhodococcal group (*R. opacus, R. jostii, R. wratislaviensis* and *R. imtechensis*) to synthesize and accumulate specific lipids of biotechnological interest (15). However, knowledge concerning sulfur metabolism and especially methionine-cysteine interconversion routes in *Rhodococcus* and other desulfurizing species, is still relatively limited.

### The Methionine-Cysteine interconversion pathways in bacteria

L-methionine and L-cysteine, the sulfur-containing amino acids responsible for *dsz* repression, are interconverted with the intermediary formation of L-homocysteine and L-cystathionine through the transsulfuration metabolic pathway. L-methionine can be converted to L-homocysteine via two possible routes (**Figure 1A**). The first requires the catalytic action of a methionine γ-lyase (MγL) for methanethiol production (16), which is then oxidized to sulfide by a methyl mercaptan oxidase (MMO) present in *Rhodococcus* strain IGTS8 (17). A direct sulfhydrylation pathway can convert sulfide to L-homocysteine, in condensation with either O-succinyl-L-homoserine (OSHS) or O-acetyl-L-homoserine (OAHS), through the catalytic action of MetZ or MetY, respectively (4, 18). A second pathway for methionine catabolism, involves the sequential formation of S-Adenosyl-L-methionine (SAM), S-Adenosyl-L-homocysteine (SAH), and L-homocysteine (4, 19–23). In the first step of the *forward* transsulfuration pathway, a γ-replacement reaction of L-cysteine and an activated L-homoserine ester (OSHS/OAHS) generates L-cystathionine, with the catalytic action of a Cystathionine γ-synthase (CγS; **Figure 1B**, Reactions M1 and M2) (24, 25). In the second forward transsulfuration step, L-cystathionine is acted upon by a Cystathionine beta-lyase (CβL) to form L-homocysteine. This in turn, can be converted to L-methionine through a methylation step or serve as the precursor for L-cysteine biosynthesis via the *reverse* transsulfuration pathway (25). Therein, a Cystathionine β-synthase (CβS)-mediated condensation of L-homocysteine with L-serine generates L-cystathionine (**Figure 1B**, Reaction C1), which is then cleaved by a Cystathionine γ-lyase (CγL) to form L-cysteine (**Figure 1B**, Reaction M5). Both key enzymes of the *reverse* transsulfuration pathway, CβS and, CγL, are pyridoxal phosphate (PLP)-dependent (26–29). This reverse transsulfuration metabolic route has been reported in mammals, yeasts, archaea and several bacteria (20, 30–34). Alternative biocatalytic reactions of CβS which generate H_2_S in the presence of cysteine, have been reported in bacterial species and eukaryotic cells. Interestingly, a condensation of cysteine with homocysteine catalyzed by CβS, can produce cystathionine (35, 36). However, H_2_S production was studied recently in knockout strains of *M. tuberculosis*, a species closely related to *R. qingshengii* IGTS8. Therein, the cysteine desulfhydrase Cds1 - but not CβS-was shown to be responsible for these reactions (37). Another pathway for L-cysteine biosynthesis, requires the O-acetyl-L-serine (OAS) sulfhydrylase, CYSK, for the condensation of sulfide and OAS (4, 38). In the opposite direction, a reaction mediated by L-cysteine desulfhydrase (CD) leads to L-cysteine degradation into sulfide, pyruvate, and ammonia (38).

**Figure 1.**
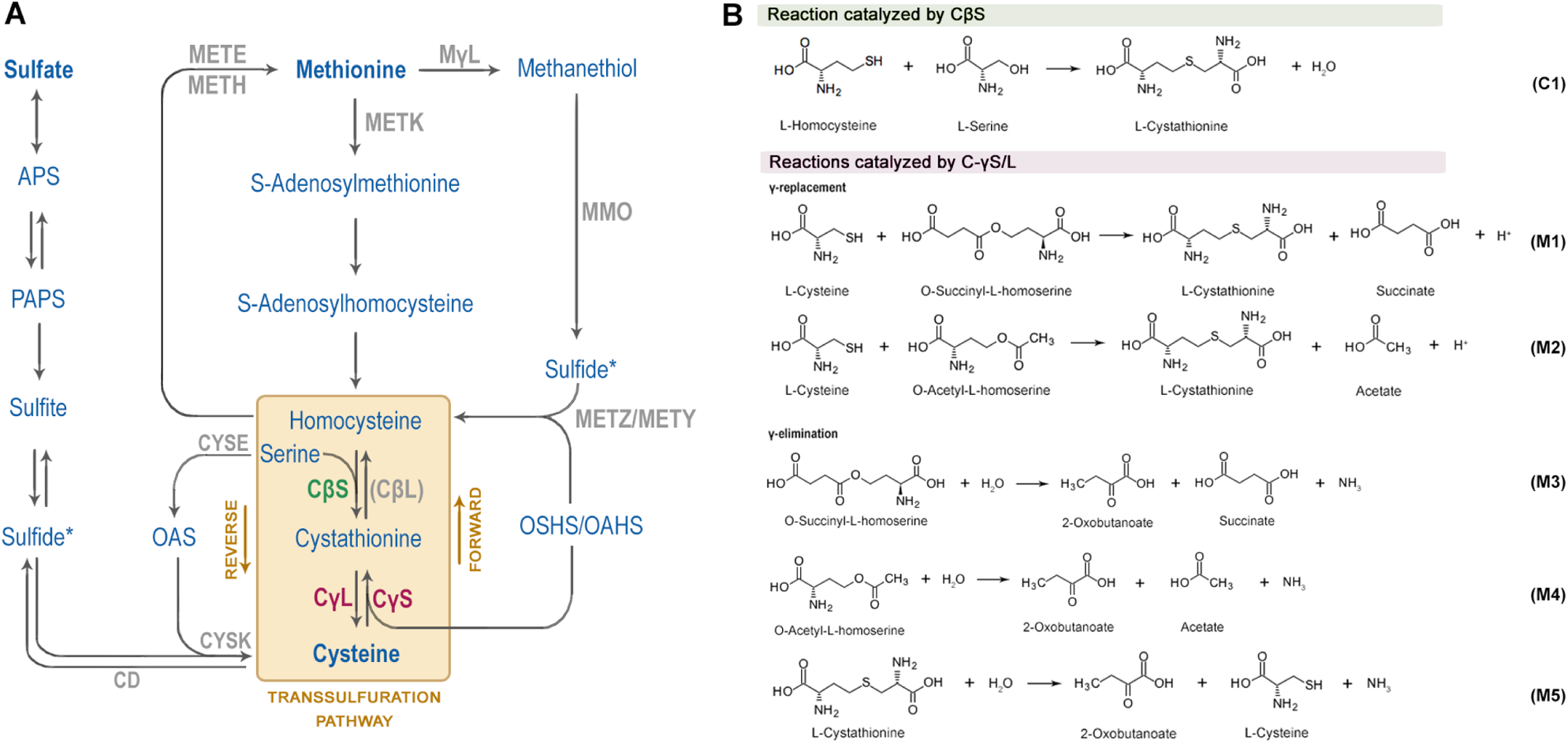
Bacterial sulfur metabolism. (A) Overview of standard Methionine and Cysteine biosynthesis and interconversion routes in bacteria as part of the sulfur assimilation pathway (APS: Adenylylsulfate, PAPS: 3’ Phosphoadenylyl sulfate, OAS: O-acetyl-L-serine, OSHS: O-succinyl-L-homoserine, OAHS: O-acetyl-L-homoserine). (B) Canonical reactions of sulfur metabolism catalyzed by CβS, METB (C-γS/L) in the *Corynebacteriales* order.

### Interconnection of transsulfuration and desulfurization pathways in Rhodococcus sp. and related species

The genome of the model biocatalyst *R. qingshengii* IGTS8, harbors genes for CβS and C-γS/L, an indication for an active reverse transsulfuration pathway. The gene product of *cbs* gene is annotated as a putative CβS Rv1077, whereas *metB* is predicted to encode a Cystathionine γ-synthase/lyase (C-γS/L). Transposon mediated disruption of the *cbs* gene has been reported in the desulfurizing strain *R. erythropolis* KA2-5-1, and it was suggested that sulfate and methionine are indirectly involved in the repression of the *dsz* phenotype (39). However, sulfur assimilation pathways and the regulation of *dsz* expression in response to different sulfur sources in desulfurizing *Rhodococcus* species, remains largely understudied *in vivo*.

Several genetic modifications have been conducted with a direct biotechnological approach, aiming to increase the efficiency of BDS rather than elucidate the underlying sulfur assimilation regulatory mechanisms. As such, most of them engineer *Escherichia coli* or *Pseudomonas* strains (40–42). However, a major limiting factor when G (−) bacteria are used as biocatalysts in a biphasic system is the mass transfer rate of DBT from the oil to the aqueous phase, that necessitates the use of co-solvents for higher efficiency (43, 44). This observation highlights the role of bacterial surface properties, such as hydrophobicity and cell wall thickness, for efficient BDS in biphasic media (45). In this regard, *Rhodococcus* biocatalysts that have the advantageous traits associated with the genus, pose as ideal candidates for genetic enhancement. However, this approach has not been favorable, especially in terms of targeted genetic modifications, owing to their extremely low amenability to genome editing (46). To date, only a few studies have used genetically engineered desulfurizing *Rhodococcus* strains, which however harbor non-stable expression vectors or randomly integrated transposon elements (39, 47–50), while none have introduced site-directed, genome-based modifications in IGTS8 or in any other *Rhodococcus* sp. desulfurizing strain.

In the present work, we generate recombinant IGTS8 biocatalysts to investigate the effects of potential gene targets on biodesulfurization activity. More specifically, we implement a precise, two-step double crossover genetic engineering approach for the deletion of two sulfur metabolism-related genes, designated *cbs* and *metB*, of *R. qingshengii* IGTS8 (51). Moreover, we provide sequence analyses of the related protein products (CβS and C-γS/L), with emphasis on highly conserved residues of the catalytic core. We present evidence that deletion of the *cbs* gene leads to derepression of BDS phenotype mostly for cells grown in the presence of sulfate, whereas BDS of the *metΒΔ* engineered strain is more prominent for methionine-grown cells. Furthermore, we report the regulatory role of both CβS and METB (C-γS/L) in *dszABC* transcription levels in response to the presence of sulfate and methionine, but not cysteine. Thus, we manage to indirectly mitigate the effect of sulfur source repression through targeted genome editing without modifying the native *dsz* operon.

## Results

### Sequence analysis of the cbs-metB genetic locus

Whole genome sequencing of *R. qingshengii* IGTS8 (51) revealed a 1386 bp ORF for *cbs* and a 1173 bp ORF for *metB*, predicted to encode for a cystathionine β-synthase – CβS, and a cystathionine γ-synthase/lyase - C-γS/L, respectively. The locus exhibits similar organization to that of KA2-5-1 strain (39) (**Figure 2A**). The gene located upstream of the *cbs-metB* locus exhibits 61% identity with *M. tuberculosis* Rv1075c, a GDSL-Like esterase (52), while the gene downstream of *metB* is predicted to encode for an L-threonine ammonia-lyase. Analysis of the upstream flanking sequence of *cbs* suggests the presence of a bacterial promoter located ∼100 bp before the *cbs* start codon (**Figure 2B**).

**Figure 2.**
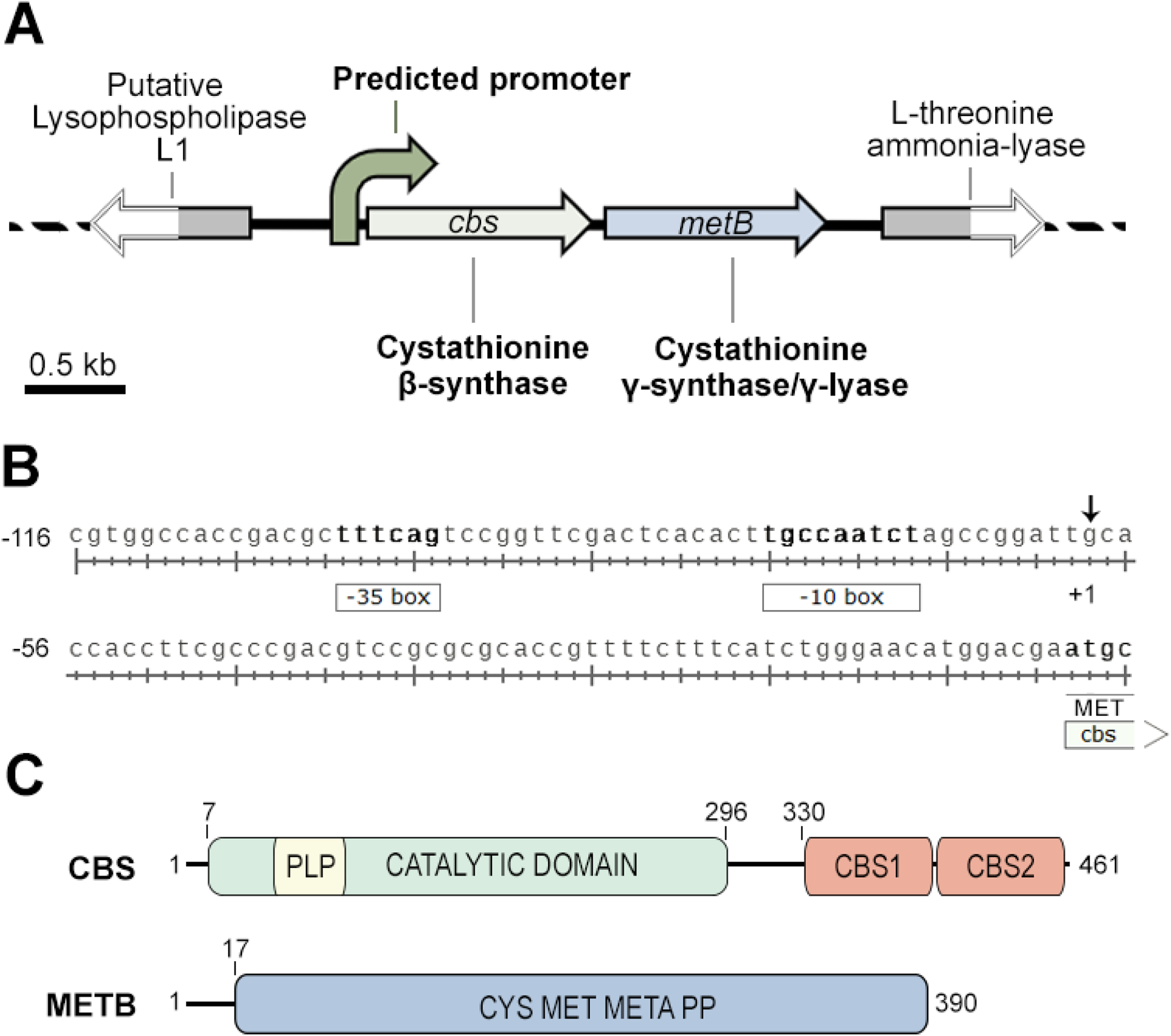
Properties of *cbs-metB* genetic loci and proteins. (A) Scheme of the *cbs-metB* gene cluster. (B) Bacterial promoter predicted sequence. −35 and −10 boxes are displayed, whereas an arrow indicates the predicted transcription initiation site (+1). (C) Schematic diagram of CβS and METB (C-γS/L) domain distribution. See main text for details.

Based on sequence homology, IGTS8 CβS consists of one N-terminal catalytic domain with the ability to bind PLP (7 - 296; pfam00291) and two C-terminal CBS regulatory motifs (CBS1, 330 - 397 and CBS2, 403 - 459; pfam00571) commonly referred to as the Bateman module (53, 54). In human and higher eukaryotes, the protein also harbors an N-terminal Heme binding domain of approximately 70 amino acid residues preceding the catalytic core domain, which has not been found in lower eukaryotes and prokaryotes (36, 55–58). METB (C-γS/L) consists of a large Cysteine/Methionine metabolism-related PLP-binding domain (pfam01053), spanning almost the entire protein length (17 - 390) (**Figure 2C**). The translated amino acid sequences of IGTS8 CβS and METB were compared to other known CβS and C-γS/L proteins, respectively. Multiple sequence alignments revealed the presence of six conserved blocks in the catalytic core of CβS and three in the C-terminal Bateman module of the protein, whereas seven blocks are identified in METB (**Figure 3**). *M. tuberculosis* CβS shows the highest similarity score to IGTS8 CβS and shares extensive homology across the entire length of the protein (99% coverage, 83% Identity). Among the other known CβS homologs, MccA from *B. subtilis*, an O-acetylserine dependent CβS, shows a 41% overall identity for the compared region (65% coverage), although this protein completely lacks the C-terminal CBS1 and CBS2 regulatory domains. The *H. sapiens* and *S. cerevisiae* counterparts show 40% and 34% similarity, respectively, throughout both the Catalytic domain and the Bateman module of the CβS protein. Residues of the catalytic cavity that interact with CβS substrates and the cofactor PLP, are extremely well conserved across the compared sequences (**Figure 3A**, blue and yellow boxes respectively), whereas alignment of the C-terminal CβS regions reveals several highly conserved residues, distributed in three blocks (**Figure 3B**).

**Figure 3.**
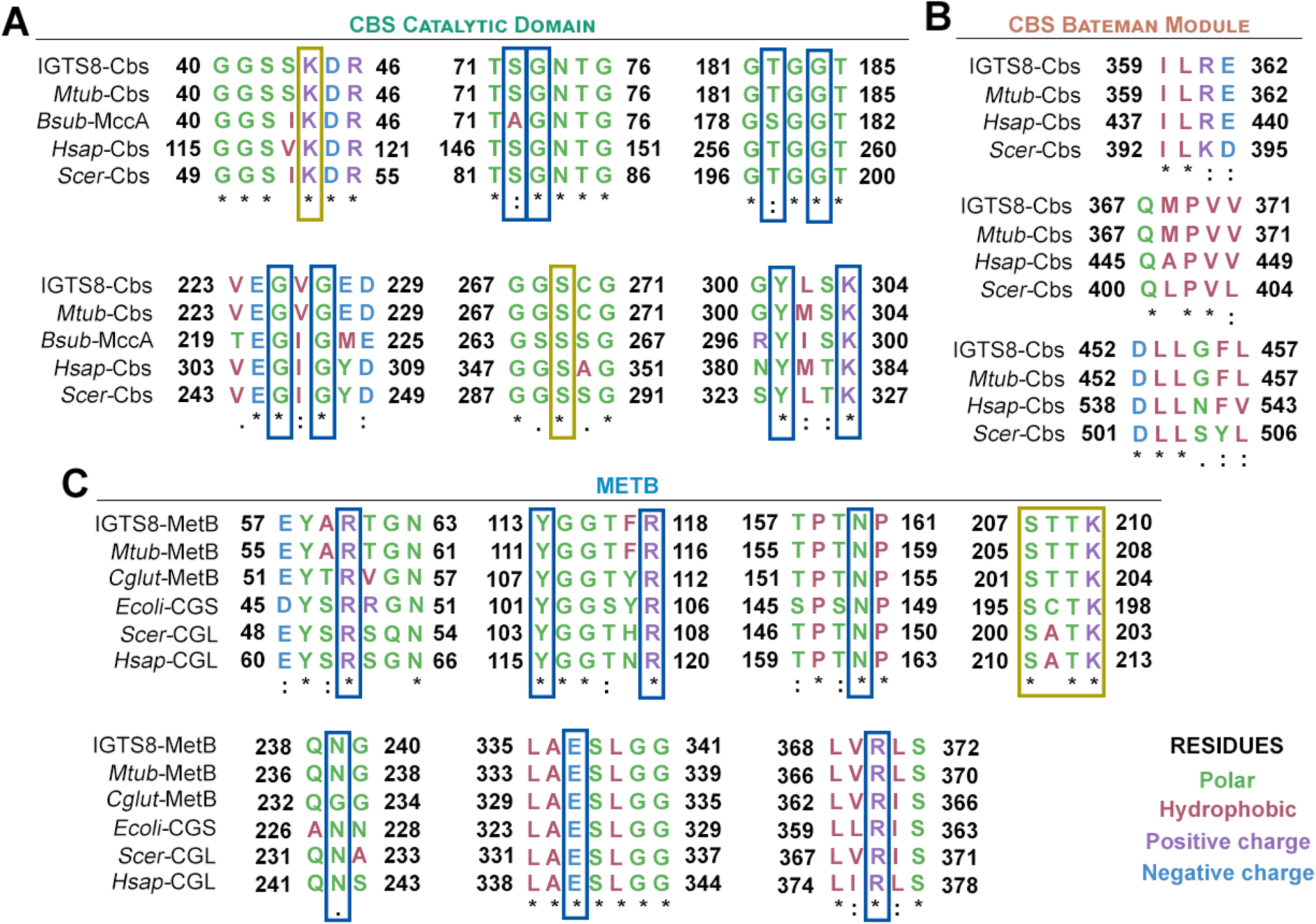
Multiple sequence alignments of CβS and C-γS/L, displaying only conserved residues configuring the active sites. (A) Comparison of *R. qingshengii* IGTS8 CβS with *M. tuberculosis* cystathionine β-synthase (Uniprot accession No: P9WP51); *B. subtilis* MccA (Uniprot accession No: O05393); Human CβS (Uniprot accession No: P35520-1); and *S. cerevisiae* CβS (Uniprot accession No: P32582). (B) Comparison of *R. qingshengii* IGTS8 METB with *M. tuberculosis* C-γS/L (Uniprot accession No: P9WGB7); *C. glutamicum* CγS (Uniprot accession No: Q79VD9); *E. coli* cystathionine γ-synthase (Uniprot accession No: P00935); *S. cerevisiae* cystathionine γ-lyase (Uniprot accession No: P31373) and Human cystathionine γ-lyase (Uniprot accession No: P32929). All multiple sequence alignments were done using ClustalO. Dashes indicate gaps introduced for alignment optimization. Asterisks (*) indicate fully conserved residues; double dots (:) denote strongly conserved residues and (.) show weakly conserved residues. Residues in yellow boxes participate in PLP-binding. Blue boxes denote residues involved in substrate binding (57, 22).

The METB (C-γS/L) multiple sequence alignment includes the *M. tuberculosis* and *C. glutamicum* METB, the cystathionine γ-synthase from *E. coli,* and the cystathionine γ-lyases from yeast and human (**Figure 3C**). *M. tuberculosis* and *C. glutamicum,* which are closely related to *R. qingshengii* IGTS8, possess homologs with the highest identity scores (73% and 65%, respectively), whereas coverage was high in all METB sequence alignments (95-99%). Notably, the CγS from the Gram-negative *E. coli* appears to have a lower similarity (42%) than the eukaryotic CγLs from *S. cerevisiae* and *H. sapiens* (49% and 47%, respectively). This observation is in line with the predicted bifunctionality of IGTS8 METB as both CγL and CγS, a unique feature that allows the synthesis of L-cysteine through L-methionine via the reverse transsulfuration pathway (25, 59, 60).

Importantly, Nucleotide Blast search within the IGTS8 genome does not reveal any additional CβS and METB homologues. In the case of CβS, tBlastN search reveals candidates with identities from 33% - 46% (IGTS8_peg5353/CysK1, IGTS8_peg6007/CysK, IGTS8_peg2567/ThrC), however with low coverage. Concerning IGTS8 METB, tBlastN search reveals four putative paralogues (IGTS8_peg1115/CTH, IGTS8_peg771/MetY, IGTS8_peg3888/MetY, IGTS8_peg5771/MetZ) again with relatively low amino acid identity (34% - 38%). Moreover, when compared to *M. tuberculosis* METB, these paralogues exhibit 35% - 36% identity and are predicted to be responsible for unrelated functions, except for IGTS8_peg1115 (CTH), proposed to also act as a putative cystathionine gamma-lyase (4).

### Effect of C source type on desulfurization activity of IGTS8

To assess the effect of different carbon sources supplementation on BDS capability and determine the corresponding preferred carbon source for *R. qingshengii* IGTS8, we collected samples from actively growing cultures at three different time-points (early-log, mid-log, and late-log phase). Wild type (wt) *Rhodococcus* cells were grown on either glucose, glycerol, or ethanol as sole carbon sources with 1 mM DMSO as sole sulfur source (**Figure 4**). The highest desulfurization activity for strain IGTS8 was obtained with the use of ethanol as a carbon source. On the contrary, utilization of glucose as a carbon source did not lead to a significant increase in attained biomass (0.12 ± 0.02 g/L) or to efficient BDS (0.30 ± 0.01 Units/mg DCW). In fact, cells did not exhibit a clear exponential growth even after 80 hours of incubation (growth rate μ_max_ and maximum biomass concentration C_max_, could not be determined). The presence of glycerol as the sole carbon source led to μ_max_ of 0.075 ± 0.01 h^−1^, C_max_ of 0.80 ± 0.06 g/L and a BDS maximum of 19.00 ± 0.04 Units/mg DCW. Comparison to the determined μ_max_ (0.081 ± 0.01 h^−1^), C_max_ (0.95 ± 0.6 g/L), and measured catalytic activity (38.0 ± 1.9 Units/mg DCW) of the same strain upon ethanol supplementation (**Figure 4A**), reveals a statistically significant difference for the compared BDS activities (**Figure 4B**), but not for the calculated growth kinetic parameters.

**Figure 4.**
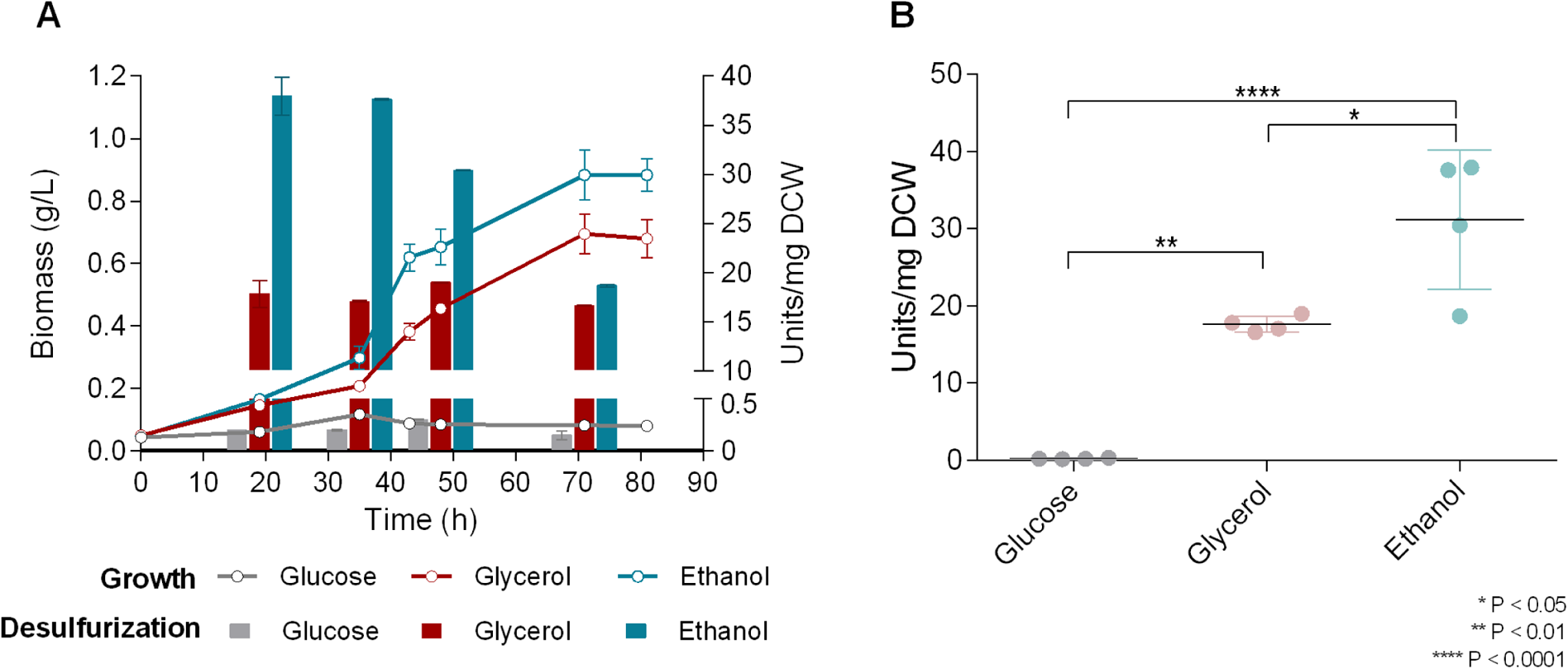
Ethanol is a preferred carbon source for *R. qingshengii* IGTS8. **(A)** Effect of different carbon sources (0.055 M Glucose, 0.110 M Glycerol, 0.165 M Ethanol) on growth (Biomass; g/L) and BDS activity (Units/mg DCW) of *R. qingshengii* IGTS8. DMSO at a concentration of 1 mM was used as the sole sulfur source. **(B)** Statistical analysis of BDS results shown in (A). One-way ANOVA with Tukey’s multiple comparison test was performed (For more details see Materials and Methods).

### Growth of knockout mutants on DBT

To study the role of Cystathionine β-synthase (CβS) and Cystathionine γ-lyase/synthase (C-γS/L, METB) in the regulation of *dsz* operon expression according to sulfur availability, scarless deletions of the corresponding genes (*cbs,* IGTS8_peg3012; and *metB,* IGTS8_peg3011) were performed with the use of the *pK18mobsacB* vector system (**Supplementary Figure S1**; For more details, please refer to the Materials and methods section). Gene deletions were verified with PCR and DNA sequencing of the PCR products. Τhe isogenic knockout strains *cbsΔ* and *metBΔ* retained their ability to grow and desulfurize on liquid minimal media in the presence of 0.1 mM DBT, without the addition of cysteine or methionine (**Supplementary Figure S2**). Ethanol was supplemented as sole carbon source (0.33 M carbon). Calculated growth rate (μ_max_) of *metBΔ* strain was marginally - but not significantly-higher than that of wt and *cbsΔ* strains, whereas neither the differences between maximum calculated biomass concentrations (C_max_), nor the produced 2-hydroxybiphenyl (2-HBP) levels (μM) were significantly different among the three strains (**Supplementary Figure S2**).

### Recombinant strains exhibit increased desulfurization activity when grown on repressive sulfur sources

We investigated the effect of *cbs* and *metB* deletions on growth and desulfurization capability of *R. qingshengii* IGTS8, comparing the isogenic *cbsΔ, metBΔ* and wt strains for: 1) maximum specific growth rates (μ_max_), 2) maximum biomass concentrations (C_max_), and 3) biodesulfurization (BDS) activities (Units/mg DCW) of resting cell suspensions harvested at three different time points (20, 45, 65 h). Ethanol was used as the sole carbon source (0.33 M carbon), whereas the sole sulfur source varied between a non-repressive (DMSO) or one of the three repressive sulfur sources (sulfate/methionine/cysteine), supplemented in two concentrations (0.1 mM or 1 mM). Interestingly, growth phase-dependent variations of BDS were observed for all strains that retained the ability to desulfurize under the influence of different medium compositions, whereas statistical analysis reveals that growth kinetic parameters of each strain were not significantly affected by the amount of sulfur source added (**Figures 5–8** and **Table S2**). The BDS phenotype of resting cells was firstly determined for cultures grown under non-repressive conditions, with the supplementation of DMSO (**Figure 5**). Comparison of C_max_ values between the wt and each of the two recombinant strains shows statistically significant differences for each supplemented concentration of DMSO (P<0.05), whereas differences between the respective μ_max_ values were non-significant. Comparison of BDS activities between different strains grown on DMSO, shows a statistically significant adverse effect of CβS depletion, whereas *metBΔ* strain is affected primarily during mid- and late-log phases (P<0.05; See also **Figure 5B** and **5C** for more details). Additionally, non-significant concentration-dependent (0.1 vs 1 mM DMSO) variations of BDS activities were observed for each strain studied.

**Figure 5.**
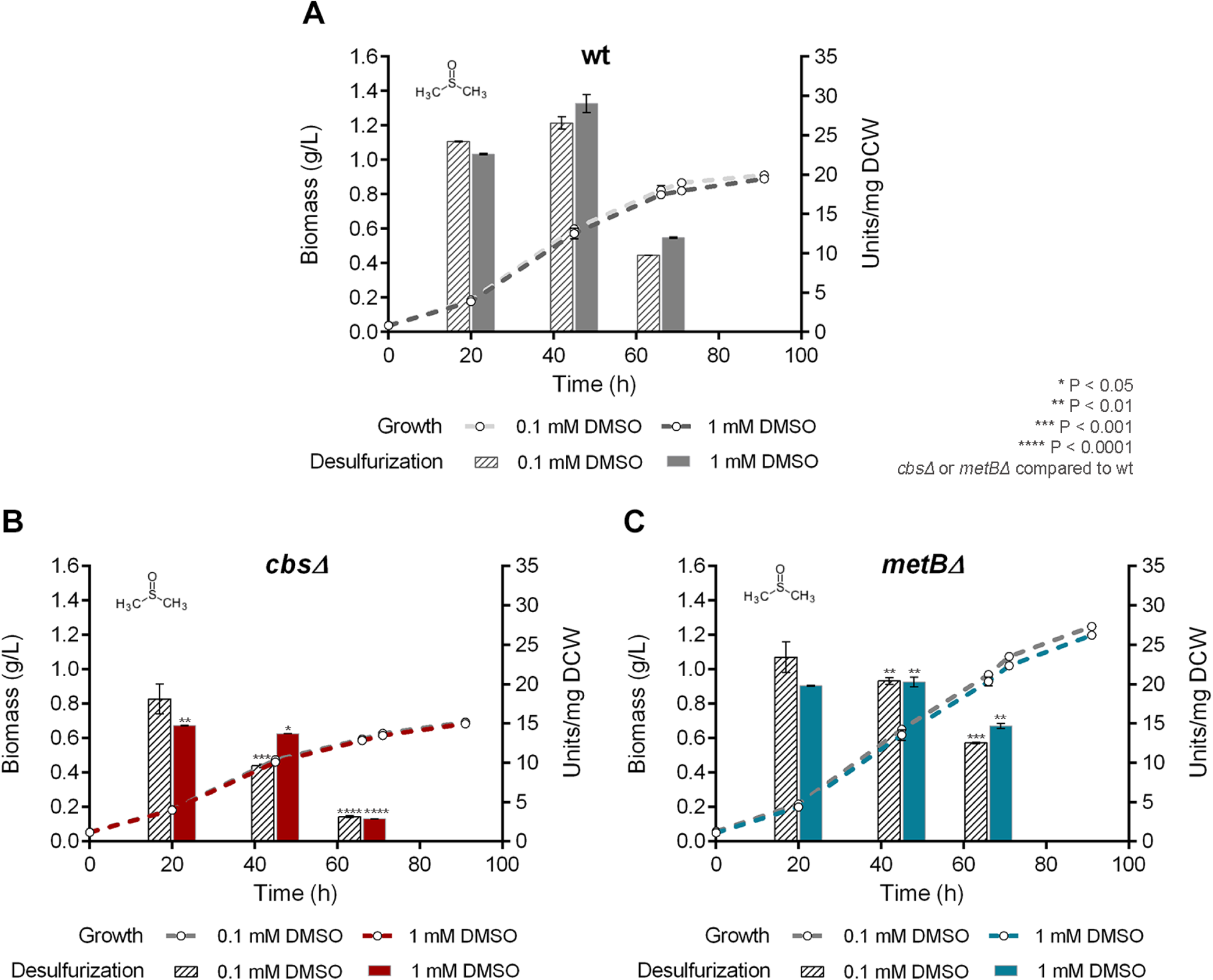
Effect of DMSO as sole sulfur source on growth and BDS activity. (A-C) Growth (Biomass; g/L) and desulfurization capability (Units/mg DCW) of wt (A), *cbsΔ* (B), and (C) *metBΔ* strains, grown on CDM in the presence of low (0.1 mM) and high (1 mM) ***DMSO*** concentrations. Ethanol (0.165 M) was supplemented as the sole carbon source in the culture medium.

To determine the effect of *cbs* and *metB* gene deletions under BDS-repressive conditions, we performed growth studies with the supplementation of sulfate, methionine, or cysteine as sole sulfur sources (**Figures 6, 7, 8** respectively and **Table S2**) followed by resting cells’ desulfurization assays. Sulfate addition in the bacterial culture efficiently represses the desulfurization phenotype of the wt strain, only when a high concentration is supplemented (1 mM; P<0.0001; **Figure 6A**). The specific growth rate of *metBΔ*, but not *cbsΔ* strain, exhibits a significant reduction in the presence of 0.1 mM sulfate (P<0.01; compared to wt). Values of calculated growth kinetic parameters are reported in **Table S2**. Deletion of *cbs* and *metB* leads to non-significant variations of calculated biomass maximum C_max_ for both sulfate concentrations, compared to wt. Notably, desulfurization is enhanced 9-fold for *cbsΔ* in the presence of 1 mM sulfate, reaching up to 15.23 ± 0.27 Units/mg DCW mid-log, compared to the wt (P<0.0001; **Figure 6A and 6B**). The METB-depleted strain exhibits a significant BDS phenotype only during the late-exponential phase for 1 mM sulfate (P<0.05; 7.49 ± 0.51 Units/mg DCW, **Figure 6C**). Moreover, concentration-dependent variations in BDS activities are validated for wt (P<0.0001; 45h and 65h), *cbsΔ* (P<0.0001 20h; P<0.01 45h), and *metBΔ* (P<0.05 45h; P<0.001 65h) strains, grown in the presence of sulfate (0.1 vs 1 mM).

**Figure 6.**
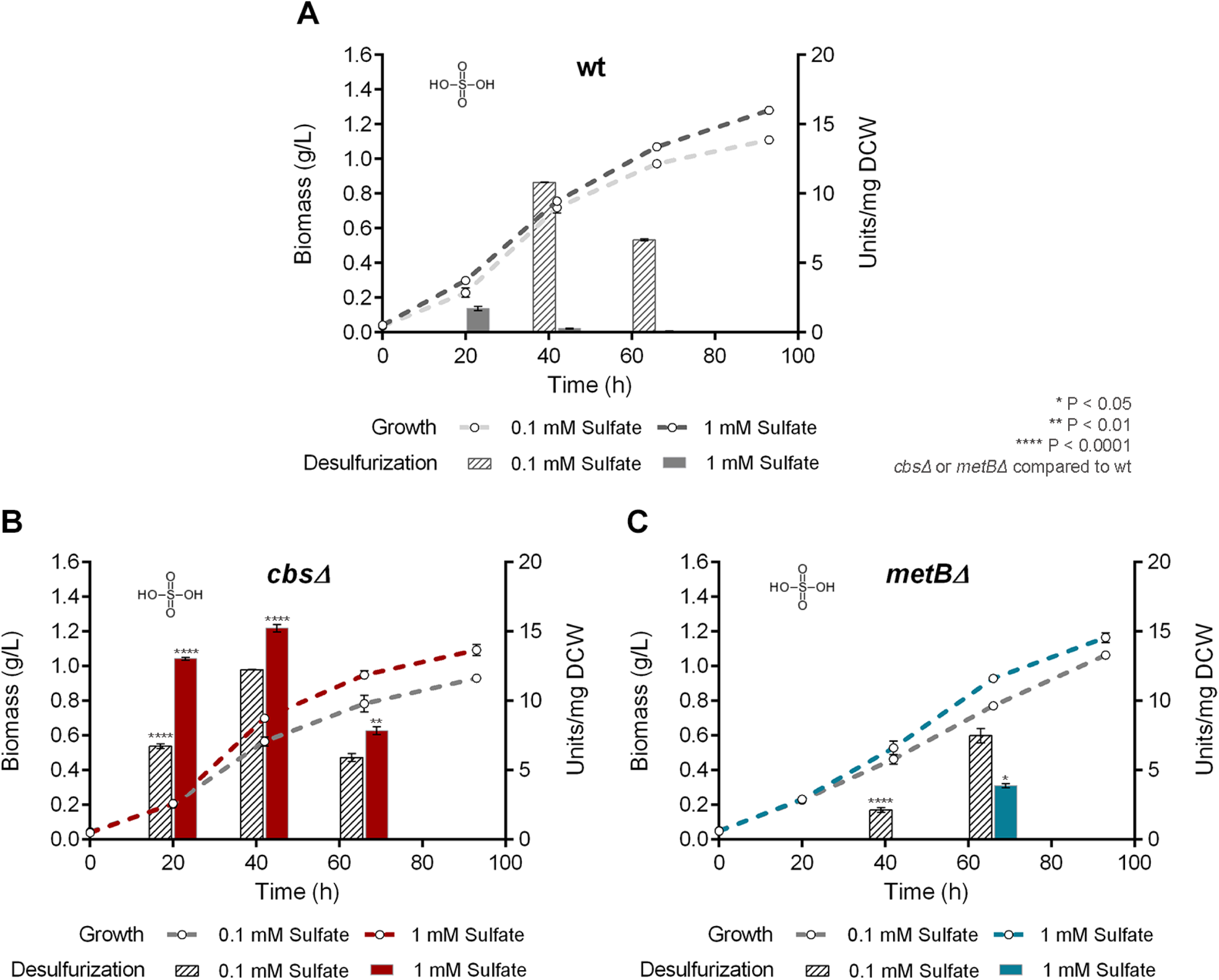
Effect of sulfate as sole sulfur source on growth and BDS activity. Growth curves (Biomass; g/L) and biodesulfurization efficiencies of resting cells (Units/mg DCW) of wt (A), *cbsΔ* (B), and *metBΔ* (C) isogenic strains, in the presence of low and high ***sulfate*** concentrations as sole sulfur sources. Ethanol (0.165 M) was supplemented as the sole carbon source in the culture medium.

**Figure 7.**
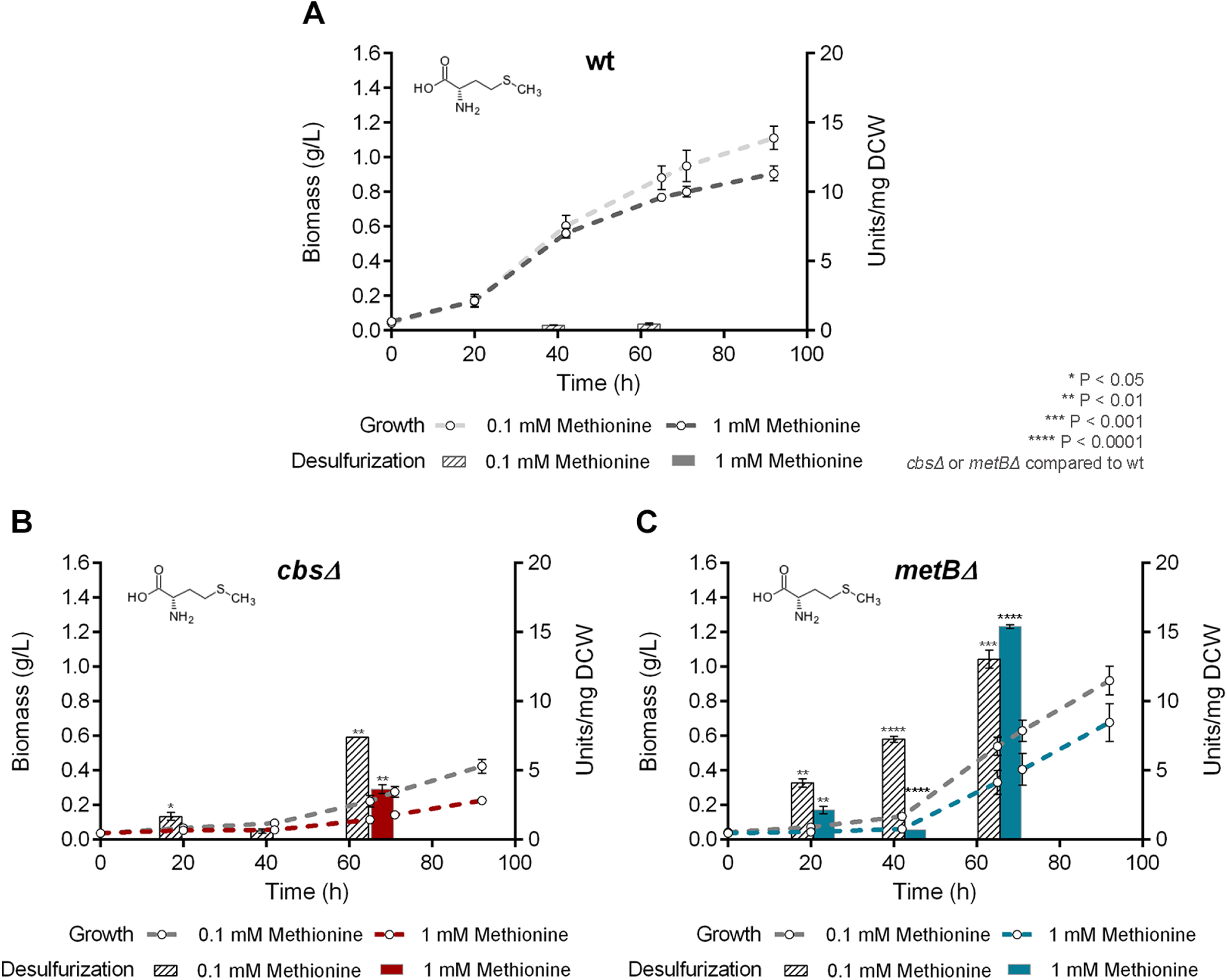
Effect of methionine as sole sulfur source on growth and BDS activity. Growth curves (Biomass; g/L) and biodesulfurization efficiencies of resting cells (Units/mg DCW) in the presence of low (0.1 mM) and high (1 mM) L-***methionine*** concentration, for wt (A), *cbsΔ* (B), and *metBΔ* (C) isogenic strains. Ethanol (0.165 M) was supplemented as the sole carbon source in the culture medium.

**Figure 8.**
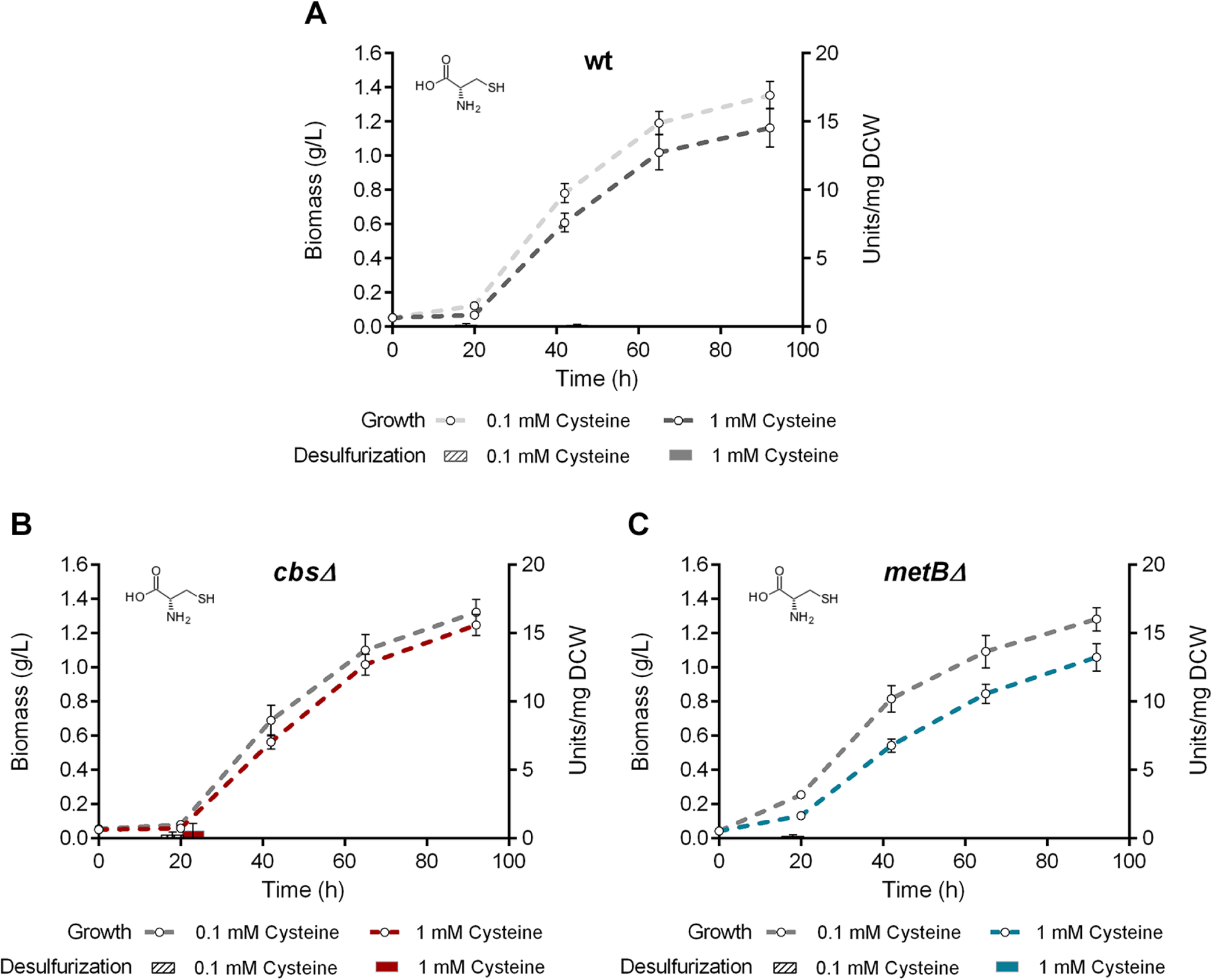
Effect of cysteine as sole sulfur source on growth and BDS activity. Growth curves (Biomass; g/L) and biodesulfurization efficiencies of resting cells (Units/mg DCW) in the presence of low (0.1 mM) and high (1 mM) L-***cysteine*** concentration, for wt (A), *cbsΔ* (B), and *metBΔ* (C) isogenic strains. Ethanol (0.165 M) was supplemented as the sole carbon source in the culture medium.

An unexpected finding is that the absence of CβS and METB seems to have a negative effect on methionine-based growth (**Figure 7 and Table S2**). Recombinant *metBΔ* exhibits lower μ_max_ values than the wt strain (P<0.05, 0.1 mM; P<0.0001, 1 mM), whereas the calculated C_max_ value for growth on 1 mM methionine appears to be significantly increased (P<0.001). Importantly, growth kinetic parameters could not be determined for *cbsΔ* strain, due to poor growth. Concerning BDS activities of strains grown on methionine, wt is completely unable to desulfurize DBT even in the presence of a low concentration (0.1 mM; 0.46 ± 0.06 Units/mg DCW; **Figure 7A**). In **Figure 7B**, the BDS phenotype of *cbsΔ* strain becomes more evident for cells harvested after 65 hours of growth on both methionine concentrations (7.34 ± 0.05 Units/mg DCW, 0.1 mM and 3.65 ± 0.31 Units/mg DCW, 1 mM; P<0.01 compared to wt). On the contrary, *metBΔ* strain exhibits remarkable desulfurization activity after 65 hours of growth on both low and high concentrations of the sulfur source (13.1 ± 0.65 Units/mg DCW, P<0.001 and 15.4 ± 0.13 Units/mg DCW, P<0.0001, respectively; **Figure 7C**). Concentration-dependent variations (0.1 vs 1 mM methionine) are also observed, when comparing BDS activities for recombinant strain *cbsΔ* (P<0.01 20h; P<0.0001 65h) and *metBΔ* (P<0.001 20h; P<0.0001 45h and 65h), and C_max_ for *metBΔ* (P<0.05).

Cysteine supplementation as the sole sulfur source in the culture medium results in complete inability of all strains (wt, *cbsΔ* and *metBΔ*) to desulfurize DBT, even in the presence of low sulfur content (**Figure 8**). However, growth is extremely efficient in all cases, as maximum biomass concentrations reach up to 1.37 ± 0.065 g/L (**Table S2**). Differences between the μ_max_ or C_max_ values of recombinant strains compared to the respective wt values, as well as concentration-dependent variations within-strain (0.1 vs 1 mM cysteine), were non-significant. We also tested the effect of cysteine supplementation at 10 mM for all strains, as cysteine is known to be toxic at high concentrations (61). Calculated C_max_ values of wt and *cbs*Δ strains, but not *metBΔ*, were significantly reduced in the presence of high exogenous cysteine concentration (10 mM), compared to lower cysteine concentrations (wt, 0.1 mM vs 10 mM: P<0.05; *cbs*Δ, 0.1 or 1 mM vs 10 mM: P<0.0001) (**Supplementary Figure S3**). Growth rates of all strains were not significantly affected by the higher cysteine concentration.

### Gene deletion of cbs or metB leads to increased transcriptional levels of dszABC desulfurization genes in the presence of selected S sources

To elucidate the effect of *cbs* and *metB* deletions on the transcriptional levels of *dszABC* desulfurization genes, as well as the regulation of *dsz, cbs* and *metB* gene expression in response to sulfur availability, we performed a series of qPCR reactions for wt, *cbsΔ* and *metBΔ* strains under repressive and non-repressive conditions. In the presence of DMSO as sole sulfur source (**Figure 9A**), *dszABC* genes are efficiently expressed regardless of *cbs* or *metB* deletions. Additionally, *cbs* and *metB* transcriptional levels do not exhibit significant changes in the presence of DMSO, compared to wt. Sulfate or methionine supplementation (**Figure 9B** and **9C**, respectively) leads to repression of *dszABC* operon expression for the wt strain, while both *cbsΔ* and *metBΔ* knockout strains exhibit increased expression levels of the desulfurization genes (*cbsΔ* sulfate: P<0.01; *metBΔ* sulfate: P<0.001; *cbsΔ* methionine: P<0.05; *metBΔ* methionine: P<0.001, compared to wt sulfate or wt methionine respectively). Moreover, under the same conditions *metB* and *cbs* gene expression appears slightly elevated, but not significantly different, for the *cbsΔ* and *metBΔ* strains, respectively, compared to wt (**Figure 9B** and **9C**). Interestingly, loss of *dszABC* transcription was observed in the presence of cysteine (**Figure 9D**), not only for wt, but also for the two knockout strains. Furthermore, *cbs* and *metB* expression levels do not exhibit significant changes between different *cbs^+^* or *metB^+^* strains grown on the same sulfur source, or for the same strain grown on different sulfur sources (**Figure 9A-C**). Overall, the results are in line with the observed sulfate- and methionine-related derepression of the desulfurization phenotype, in response to *cbs* and *metB* deletions.

**Figure 9.**
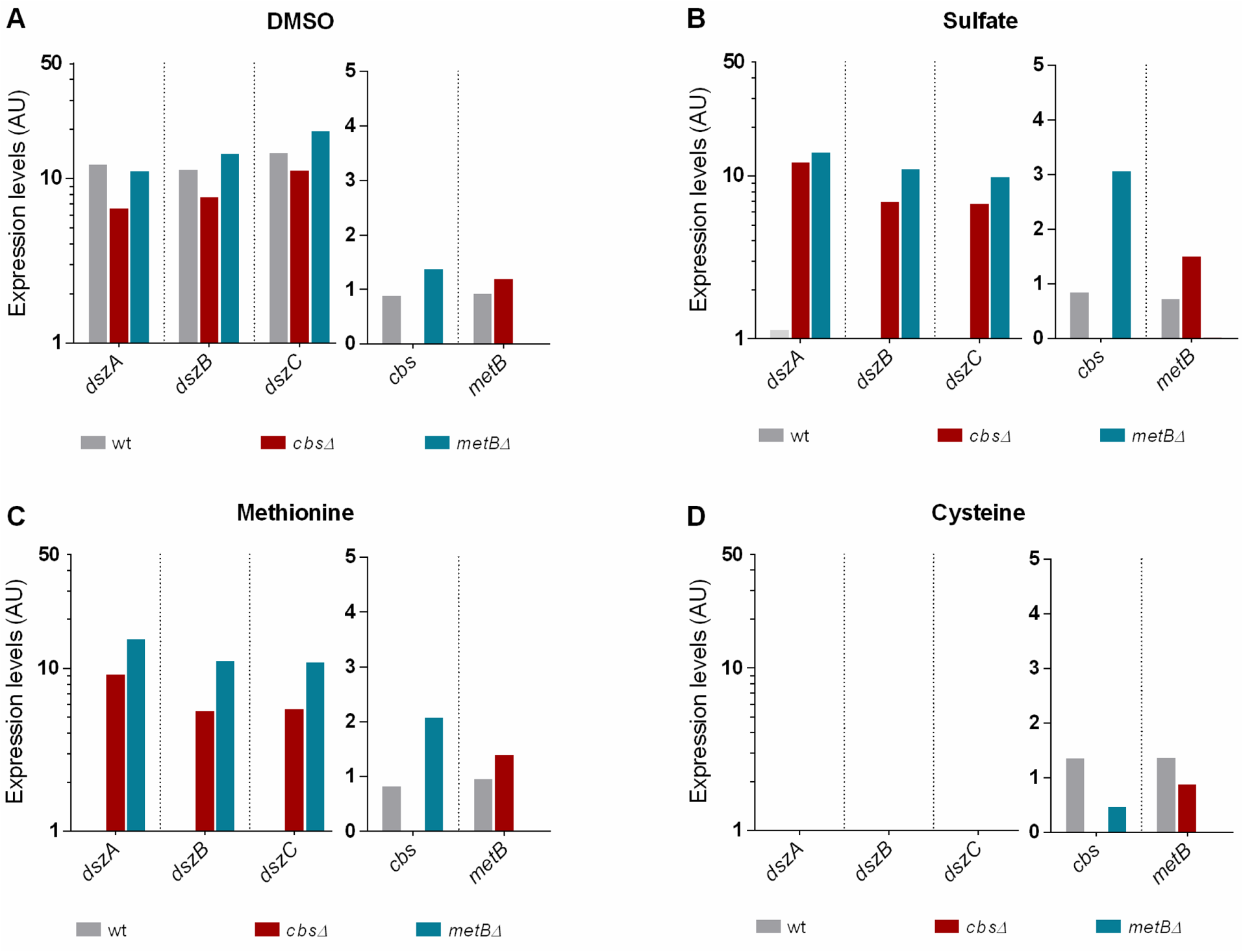
CβS and METB are critical for desulfurization genes (*dszABC*) expression in the presence of sulfate and methionine. Comparison of *dszA, dszB, dszC, cbs* and *metB* transcriptional levels for wt, *cbsΔ* and *metBΔ* isogenic strains, grown on 1 mM (A) DMSO, (B) Sulfate, (C) Methionine, or (D) Cysteine. Samples were collected from mid-log phase cultures (AU: Arbitrary Units; Relative expression levels compared to the calibrator sample. Logarithmic scale is used for *dszABC*. For details see Materials and Methods). Ethanol (0.165 M) was supplemented as the sole carbon source in the culture medium.

## Discussion

Even though certain aspects of sulfur metabolism are generally well characterized in other Gram-positive bacteria, such as *B. subtilis,* the regulation of sulfur-assimilation-related gene expression remains unclear in *Rhodococcal* desulfurizing species. This is a curiously paradoxical situation, given that *R. qingshengii* IGTS8 is the most extensively studied biocatalyst for industrial biodesulfurization applications. The main reason for this, is partly the challenging nature of genetic engineering for the actinomycete genus *Rhodococcus*, due to its high GC-content and prohibitively low homologous recombination efficiencies (46, 62). To our knowledge, no other studies have reported targeted, genome-based manipulations in desulfurizing *Rhodococci*. Contrastingly, plasmid-based modifications have been commonly used, which, however, are less preferred for industrial-scale applications as they exhibit a lower degree of genetic stability. Other approaches to date only include the *in silico* modeling of sulfur assimilation and the most recent proteomics and metabolomics analyses in strain IGTS8 (4, 12).

In the present work, we performed targeted and precise editing of the *R. qingshengii* IGTS8 genome for the first time, generating recombinant biocatalysts that harbor gene deletions of the two enzymes predicted to be involved in the reverse transsulfuration pathway. Importantly, primary amino acid sequence analyses of IGTS8 CβS and C-γS/L (METB), suggested the presence of highly conserved residue blocks that participate in active site configuration, binding of substrates and of the PLP cofactor. The high degree of similarity between the two IGTS8 enzymes and their respective counterparts found in the closely related species (63), *M. tuberculosis* (83% Identity for CβS, 73% Identity for METB), suggests a conserved function for these two proteins as Cystathionine β-synthase and Cystathionine γ-lyase, respectively. This result is in line with previous reports suggesting the existence of an operational reverse transsulfuration pathway for the genus *Rhodococcus* (4, 39). In accordance with our sequence analyses and multiple alignment results, the sulfur assimilation model proposed by Hirschler et al. (4), also identified CβS as a cystathionine β-synthase and METB as a cystathionine γ-lyase (CγL). Notably, annotation of IGTS8 genome (4) also predicts the existence of a second CγL (EC: 4.4.1.1/CTH), which however exhibits relatively low amino acid identity compared to METB of *M. tuberculosis* and IGTS8 (35% and 38%, respectively). Therefore, the possibility for an auxiliary or cryptic role of CTH in cysteine biosynthesis from cystathionine cannot be excluded.

To perform growth and desulfurization assays with the use of a single carbon source, we compared the effects of glucose, glycerol, and ethanol supplemented at 0.33 M carbon. To our knowledge, ethanol superiority as a carbon source has been verified for *R. erythropolis* KA2-5-1 (64), for transformants of *R. qingshengii* CW25, but only in comparison to glucose (50) and very recently for *R. jostii* RHA1, in comparison to glucose, n-octane, and 1-butanol (14). In the current study, ethanol is validated as the preferred carbon source for desulfurization activity of IGTS8 and is thus employed for the comparisons between wt and recombinant strains, grown in the presence of five different sulfur sources (DBT, DMSO, sulfate, methionine, cysteine). Importantly, knockout strains *cbsΔ* and *metBΔ* do not require supplemental cysteine or methionine, as evidenced by their ability to grow in the presence of DBT, DMSO, and sulfate as sole sulfur sources. The prototrophic nature of recombinant strains suggests the existence of alternative operating routes for sulfur-containing amino acid biosynthesis. Hirschler et al. (4) suggested that sulfate addition in the culture medium probably necessitates reverse transsulfuration metabolic reactions as the primary route for cysteine biosynthesis from methionine, and that the CysK-dependent alternative route for cysteine biosynthesis seems to be preferred under sulfate-starvation conditions (DBT cultures). The latter route is likely operating as a secondary pathway when sulfate is supplemented as sole sulfur source. According to the same study (4), protein levels of CβS and MetB were slightly higher, but not significantly different between the DBT and sulfate cultures. Cysteine biosynthesis via the CysK-mediated direct sulfhydrylation pathway - or other operational cysteine biosynthesis routes - could partly compensate for the reduced cysteine supply in the absence of CβS or METB in our growth and BDS studies. On the contrary, the CTH-mediated pathway for cysteine production from cystathionine could only explain the non-essentiality of MetB, but not of CβS.

In our work, growth studies reveal that DBT supplementation as sole sulfur source (0.1 mM), favors the growth rate of *metΒΔ* only marginally, compared to other strains. Contrastingly, the *metBΔ* strain exhibits significantly slower growth rates in the presence of sulfate (0.1 mM) and methionine (0.1 and 1 mM) as sole sulfur sources, but maximum biomass concentration of the recombinant is significantly elevated in the presence of DMSO (0.1 and 1 mM) and methionine (1 mM). Concerning the *cbsΔ* strain, kinetic parameters for methionine-based growth could not be determined, owing to an extremely slow growth rate. This observation is in line with the low growth yield reported for the transposon-disrupted *cbs* strain of *R. erythropolis* KA2-5-1, in the presence of 5 mM methionine (39). Although CβS and METB are non-essential, their depletion could lead to diverse effects including lower cysteine availability and/or redirection of metabolic precursors toward competing pathways. It is possible that depletion of the reverse transsulfuration pathway combined with methionine supplementation as sole sulfur source, leads to accumulation of intermediary toxic metabolites such as homocysteine or cystathionine, that could potentially inhibit growth (61, 65). Importantly, cysteine supplementation at 0.1 and 1 mM, allows or even promotes growth for the wt and both knockout strains, based on the calculated C_max_ values. Notwithstanding, a high concentration of the amino acid (10 mM) has significant adverse effects on biomass concentration maxima of wt and *cbs*Δ strains, but not of *metB*Δ. Cysteine-mediated toxicity at high concentrations, has been reported to inhibit yeast growth (61). The fact that *metB*Δ strain remains mostly insensitive to cysteine abundancy in the culture medium, concomitant with slightly elevated (although non-significant) μ_max_ values for lower cysteine content, could indicate a relatively more prominent cysteine deficiency due to the absence of METB.

Overall, our desulfurization results reveal that activity of the wt and recombinant IGTS8 strains, is largely affected by type and concentration of the sulfur source available, and in certain occasions by the growth phase of the culture. Comparison of BDS activities for different strains grown on DMSO shows a negative effect of CβS - and to a lesser extent METB - depletion. This finding is surprising, as DMSO is used widely as a non-repressive sulfur source (66, 67), however, it is possible that metabolic alterations associated with reverse transsulfuration enzyme depletion, in combination with DMSO-induced stress (68, 69), produces this adverse effect on BDS activities of recombinant strains. Sulfate addition in the bacterial culture at 1 mM, efficiently represses the desulfurization phenotype of wt, but importantly, does not affect the desulfurization activity for the *cbsΔ* strain. Deletion of *metB* also promotes a significant BDS phenotype for sulfate-based growth, but a remarkable desulfurization activity (up to 33-fold increase compared to the wt) is observed for methionine-grown *metBΔ* cells, regardless of the concentration supplemented. On the other hand, growth on cysteine completely represses the desulfurization phenotype for all strains even in the presence of low sulfur content. This is in line with the results reported by Tanaka et al. (39) for *R. erythropolis* KA2-5-1, as sulfate and methionine did not seem to be directly involved in the repression system, contrastingly to cysteine. Therefore, cysteine supplementation even at 0.1 mM, is probably an impeding factor for DBT biodesulfurization regardless of the reverse transsulfuration pathway functionality.

Concerning growth phase-dependent variations of BDS activity, in most cases, maximum values are attained mid-log and minimum BDS activity is observed for late-exponential phase cultures. Exceptions to the rule are the *metBΔ* strain supplemented with sulfate, and both recombinant strains grown on methionine, as they exhibit maximum BDS values after prolonged growth (65 h). Importantly, certain BDS activity variations are described for recombinant cells grown on different sulfate or methionine concentrations, but not for DMSO or cysteine. For example, strain *cbsΔ* prefers a higher sulfate and a lower methionine content, whereas *metBΔ* strain prefers the higher methionine concentration only for late-log harvested cells.

A possible explanation for the slightly different BDS activities exhibited by recombinant strains grown at different sulfur types and concentrations, involves alterations in the levels of CβS and METB substrates, i.e., homocysteine, serine for *cbsΔ*, and cystathionine for *metBΔ* strain. Precursor metabolite accumulation and/or redirection toward competing pathways, could signal the differential regulation of alternative sulfur assimilation routes. Some pathway examples are the MetE/MetH-mediated conversion of homocysteine to methionine, or cysteine production from serine with the intermediary formation of OAS *via* the CysE-CysK bienzyme complex (70, 71; Figure 1). Following a pathway alternative to the typical MetK-dependent, methionine can be converted to methanethiol and eventually sulfide, which in turn can either enter the MetZ/MetY-dependent routes for homocysteine production or get converted to cysteine *via* CysK. Supplemented DMSO can be reduced to dimethyl-sulfide (DMS), and subsequently converted to methanethiol, which in turn is oxidized to generate sulfide (72). Sulfate supplementation as sole sulfur source can also eventually lead to sulfide production via a four-step process, with the intermediary formation of sulfite. The availability of sulfate is known to stimulate divergent routes for sulfate/sulfite reduction, while the latter serves as a metabolic branching point (4). Changes in pathway fluxes of recombinant strains might affect BDS activity indirectly, or exert their effects at the post-transcriptional level, given that *dsz* expression levels do not seem to differ significantly for the two knockout strains in the presence of sulfate or methionine. The expression levels or the activities of proteins involved indirectly in BDS, might also be affected by alterations in the metabolic profile of recombinants. Such examples are the oxidoreductase DszD and relevant cofactor levels (NADH, FMN), or proteins involved in the import of DBT. Notably, cysteine is known to regulate the expression of genes involved in sulfur assimilation, *via* modulating CysK complex formation with the acetyltransferase CysE or the transcription factor CymR, as partners (Discussed in more detail below) (71).

As evidenced by transcript level comparison for wt, *cbsΔ* and *metBΔ* strains, CβS and METB exert an effect on *dszABC* gene expression, in response to the supplementation of different sulfur sources. Under physiological conditions, the two reverse transsulfuration enzymes regulate cysteine biosynthesis and likely promote a slight increase of the free cysteine pool, when either sulfate or methionine is used as the sole sulfur source. Deletions of the two genes might lead to reduction - but not depletion - of intracellular cysteine levels, promoting a global fine-tuning of sulfur-starvation-induced proteins expression. This in turn, could eventually allow for *dszABC* efficient expression in *cbsΔ* and *metBΔ* strains, under sulfate- or methionine-rich conditions, given that sulfur-assimilation-gene expression is widely modulated in response to sulfur source availability (**Figure 10**) (4, 73). This hypothesis is in line with the complete lack of biodesulfurization activity and the non-detectable *dsz* genes expression that was observed for the wt and knockout strains, in the presence of exogenously supplemented cysteine as the sole sulfur source. The mechanistic details of cysteine effects on *dsz* repression have not been elucidated in *R. qingshengii* IGTS8, therefore we cannot exclude a direct binding of the metabolite to a homologue of the *dsz* operon repressor WhiB1 (8). However, it is possible that cysteine promotes the CysK-CymR complex formation, via inhibition of the CysE-CysK bienzyme complex (70, 71). Importantly, homologues of CysK, CysE, and CymR are harbored within the IGTS8 genome (Blast analysis; Data not shown). Once activated, CymR (74, 75) - or some other global regulator - could have a direct repressive effect on the *dsz* operon or modulate the regulators DszGR and WhiB1 (6–8). In addition to DszGR, the general DNA binding protein IHF was also shown to be necessary for *P_dsz_*promoter activity, but its role has not been validated via knockout of the *mihf* gene in *R. qingshengii* IGTS8 or *Gordonia* sp. (9). Interestingly, an initial study postulated the existence of a ligand binding site in DszGR for sodium sulfate, leading to a change in the structure of the protein *in vitro*. This approach, however, did not conclusively show a direct role for sulfate in *dsz* operon repression for *Rhodococcus* strain IGTS8 (7). Based on our findings in *Rhodococcus* strain IGTS8, *dszABC* gene expression is no longer repressed in the presence of 1 mM sulfate once CβS or MetB are depleted. However, it is also possible that regulation differs slightly from one genus (*Gordonia*) to another (*Rhodococcus*), given that regulatory sequences of the *P_dsz_* promoter regions are only partially conserved (10). Moreover, we cannot exclude the presence of additional regulators within the IGTS8 genome that could have dominant or synergistic effects on *dsz* operon regulation *in vivo*, such as the CymR homologue described in detail above, or the global CysB regulator (76). Taken together, several mechanistic details remain to be elucidated, offering valuable information on BDS regulation.

**Figure 10.**
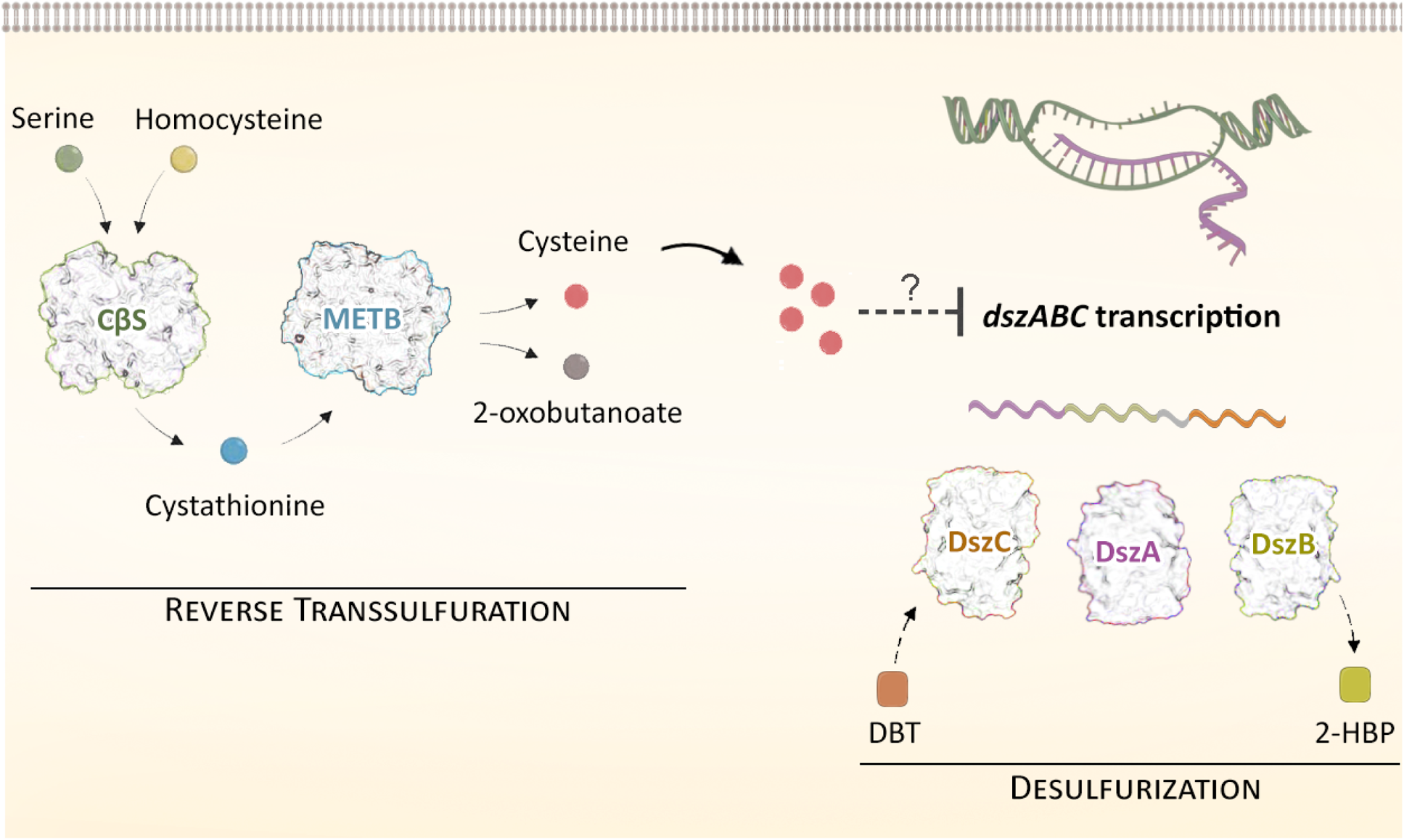
Proposed model illustrating the role of CβS and MetB (acting as CγL) in the regulation of desulfurization phenotype in *Rhodococcus qingshengii* IGTS8. Sulfate or methionine addition in the culture media, most likely necessitates reverse transsulfuration metabolic reactions as the primary route for cysteine biosynthesis. Fine-tuning of sulfur assimilation via intracellular cysteine levels is a common theme in bacterial species, where it seems to have evolved as a cellular mechanism to control gene expression appropriately, based on the available sulfur source type and abundancy. Narrow alterations in the free cysteine pool are suspected to exert an effect (directly or indirectly) on *dszABC* gene expression, leading to lack of biodesulfurization activity. Gene deletions of *cbs* or *metB*, abolish *dsz* repression in the presence of selected sulfur sources, such as sulfate and methionine, thus leading to detectable transcript levels and biodesulfurization activity.

In conclusion, our approach provides significant insights on the metabolic engineering of sulfur metabolism in *Rhodococcus qingshengii* IGTS8 without manipulation of the 4S pathway genes and reveals an important role for both CβS and METB in the global regulation of Dsz-mediated sulfur acquisition from organosulfur compounds such as DBT. From our own observations and the available literature data, we propose the involvement of both enzymes in the reverse transsulfuration pathway of *Rhodococcus qingshengii* IGTS8, we highlight an important - yet unexplored - role for cysteine in *dsz* gene expression and BDS phenotype, and we validate the necessity of intact *cbs* and *metB* loci for the orchestration of *dsz-*mediated sulfur assimilation, in response to sulfur source availability.

## Materials and Methods

### Strains, growth conditions, and plasmids

The bacterial strains and plasmids used in this study are listed in Table 1. *Rhodococcus qingshengii* IGTS8 was obtained from ATCC (53968; Former names of the strain: *R. rhodochrous, R. erythropolis*). *Escherichia coli* DH5a and S17-1 strains were used for cloning and conjugation purposes, respectively. *Rhodococcus qingshengii* strains were routinely grown in Luria-Bertani Peptone (LBP) broth (1% w/v Bactopeptone, 0.5% w/v Yeast extract, and 1% w/v NaCl) at 30°C with shaking (180-200 rpm), or on LBP agar plates at 30°C. *E. coli* strains were grown in LB medium (1% w/v Bactotryptone, 0.5% w/v Yeast extract, and 1% w/v NaCl) at 37°C with shaking (180-200 rpm) or on LB agar plates at 37°C. Kanamycin (50 μg/ml) was used for plasmid selection in *E. coli.* Kanamycin (200 μg/ml) and nalidixic acid (10 μg/ml) were used to select *R. qingshengii* transconjugants in the culture media. Counter-selection was performed on no-salt LBP (NSLBP) plates with 10% (w/v) sucrose.

**Table 1.**
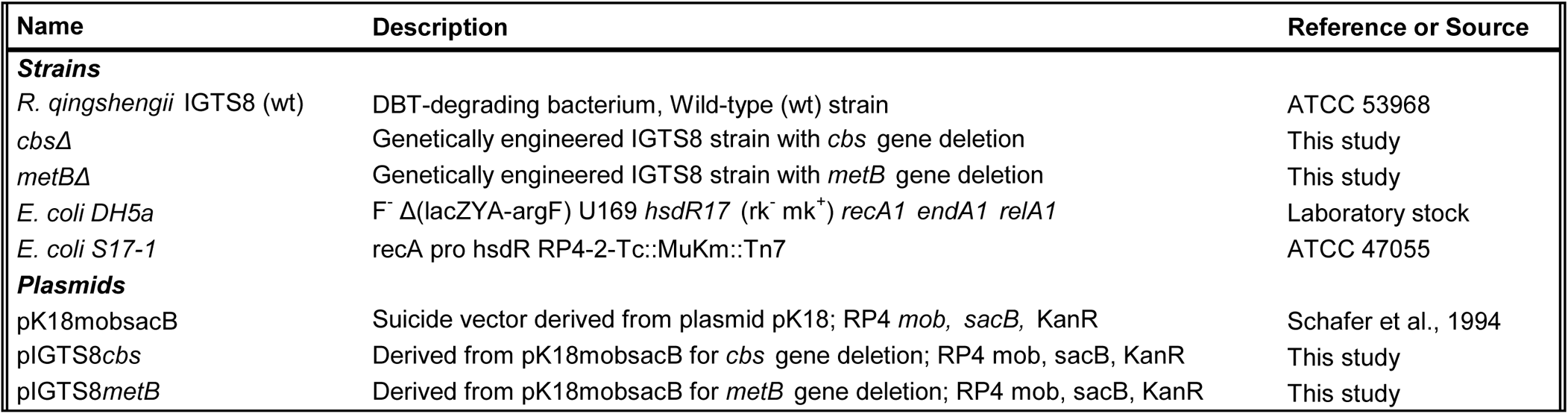
Bacterial strains and plasmids used in this study.

For biodesulfurization studies, *R. qingshengii* wild-type (wt) and recombinant strains were grown on a sulfur-free chemically defined medium (CDM) containing 3.8 g NaH_2_PO_4_·H_2_O, 3.25 g Na_2_HPO_4_·7H_2_O, 0.8 g NH_4_Cl, 0.325 g MgCl_2_·6H_2_O, 0.03 g CaCl_2_·2H_2_O, 8.5 g NaCl, 0.5 g KCl, 1 ml metal Solution, and 1 ml of vitamin solution in 1 L of distilled water (pH 7.0). The metal solution composed of (per L of distilled water): Na_2_-EDTA, 5.2 g; FeCl_2_·4H_2_O, 3 mg; H_3_BO_3_, 30 mg; MnCl_2_·4H_2_O, 100 mg; CoCl_2_·6H_2_O, 190 mg; NiCl_2_·6H_2_O, 24 mg; CuCl_2_, 0.2 mg; ZnCl_2_, 0.5 mg; Na_2_MoO_4_·2H_2_O, 36 mg; Na_2_WO_4_·2H_2_O, 8 mg; and Na_2_SeO_3_·5H_2_O, 6 mg. The vitamin solution contained (per L of distilled water) calcium pantothenate, 50 mg; nicotinic acid, 100 mg; *p-*aminobenzoic acid, 40 mg; and pyridoxal hydrochloride, 150 mg. CDM was supplemented with dibenzothiophene (DBT) (0.1 mM) or with dimethyl sulfoxide (DMSO), sulfate, L-methionine, L-cysteine as the sole sulfur source (0.1 or 1 mM) and 0.165 M ethanol, 0.055 M glucose, or 0.110 M glycerol as carbon sources (0.33 M carbon), depending on the experiment. *pK18mobsacB* (Life Science Market, Europe) was used as a cloning and mobilization vector.

### Enzymes and chemicals

All restriction enzymes were purchased from TaKaRa Bio or Minotech (Lab Supplies Scientific SA, Hellas). Chemicals were purchased from Sigma-Aldrich (Kappa Lab SA, Hellas) and AppliChem (Bioline Scientific SA, Hellas). Conventional and high-fidelity PCR amplifications were performed using KAPA Taq DNA and Kapa HiFi polymerases, respectively (Kapa Biosystems, Roche Diagnostics, Hellas). All oligonucleotides were purchased from Eurofins Genomics (Vienna, Austria) and are listed in supplementary Table S1.

### Construction of knockout strains

The genomic DNA of *Rhodococcus* strain IGTS8 was isolated using the NucleoSpin Tissue DNA Extraction kit (Macherey-Nagel, Lab Supplies Scientific SA, Hellas) according to the manufacturer’s instructions. The online software BPROM was used for bacterial promoter prediction (http://www.softberry.com/cgi-bin/programs/gfindb/bprom.pl). Unmarked, precise gene deletions of Cystathionine β-synthase (*cbs*; IGTS8_peg3012) or Cystathionine γ-lyase/synthase (*metB*; IGTS8_peg3011) were created using a two-step allelic exchange protocol (77). Upstream and downstream flanking regions of the *cbs* gene of strain IGTS8 were amplified and cloned into the pK18mobsacB vector (78), using the primer pairs *cbsUp-F/Up-R* and *cbsDown-F/Down-R*, respectively, yielding plasmid pIGTS8*cbs*. Similarly, for the flanking regions of *metB* gene, primer pairs *metBUp-F/Up-R* and *metBDown-F/Down-R* were used to construct plasmid pIGTS8*metB*. Plasmid preparation and DNA gel extraction were performed using the Nucleospin Plasmid kit and the Nucleospin Extract II kit (Macherey-Nagel, Lab Supplies Scientific SA, Hellas). *E. coli* S17-1 competent cells were transformed with each of the modified plasmids. *R. qingshengii* IGTS8 knockouts were created after conjugation (79) with *E. coli* S17-1 transformants, using a two-step homologous recombination (HR) process (Supplementary Figure S1). Following the first crossover event, sucrose-sensitive and kanamycin-resistant IGTS8 transconjugants were grown in LB overnight with shaking (180 rpm), to induce the second HR event. Recombinant strains were grown on selective media containing 10% (w/v) sucrose and tested for kanamycin sensitivity, to remove incomplete crossover events. Gene deletions *cbsΔ* and *metBΔ* were identified with PCR and confirmed by DNA sequencing of the PCR products (Eurofins-Genomics, Vienna, Austria), using external primer pairs *cbs-5F-check/cbs-3R-check* and *metB-5F-check/metB-3R-check*, for *cbsΔ* and *metBΔ* respectively.

### Growth and desulfurization assays

For inoculum preparation, cells were harvested from LBP plates in Ringer’s buffer pH=7.0, centrifuged at 3.000 rpm for 10 min and medium was discarded. The wash process was repeated twice. Pellet was resuspended in 0.5 ml of the same buffer, and necessary dilutions were performed for OD_600_ measurements and inoculum standardization. Biomass concentration, expressed as Dry Cell Weight (DCW), was estimated by measurement of optical density at 600 nm with a Multiskan GO Microplate Spectrophotometer (Thermo Fisher Scientific, Waltham, MA USA), and calculations were based on an established calibration curve. Initial inoculum concentration was determined, and an initial biomass concentration of 0.045-0.055 g/L was applied for each growth condition.

For growth studies and resting cells biodesulfurization assays (Figures 4–8), wt and recombinant strains were inoculated as described above and grown in CDM under different carbon and sulfur source types and concentrations (described in the respective figure legends). Growth took place in 96-well cell culture plates, with the use of 17 identical well-cultures per strain and condition (F-bottom; Greiner Bio-One, Fischer Scientific, US) with 150 μl working volume in thermostated plate-shakers at 30 °C and 600 rpm (PST-60HL, BioSan, Pegasus Analytical SA, Hellas). Biomass concentration at all time points was calculated as described above.

For growth and desulfurization studies in the presence of DBT (Supplementary Figure S2), *R. qingshengii* IGTS8 wt, *cbsΔ,* and *metBΔ* strains were grown in CDM with the supplementation of 0.1 mM DBT (from a 100 mM ethanol stock) as sole sulfur source, and ethanol at a final concentration of 0.165 M, as sole carbon source. Three identical 100 ml flasks with 20 ml working volume were used for each strain.

For modelling microbial growth, a simple unstructured logistic kinetic model was employed

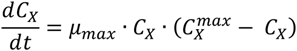

where, C_X_, is the biomass concentration in g/L, C_X_^max^, is the final biomass concentration (g/L), and μ_max_ is the apparent maximum specific growth rate (h^−1^). A nonlinear regression (SigmaPlot, version 12) routine was used to determine the model parameters, C^max^ and μ_max_, for each experimental data set (C_X_ vs t) using the integrated form of the above equation. The minimization of the sum of squared residuals was used as the convergence criterion.

Resting-cells biodesulfurization assays were performed simultaneously with the growth studies, using the cells of the corresponding 96-well plates cultures. The content of 2 to 4 identical well-cultures was harvested at early-, mid- and late-exponential phase, centrifuged at 3.000 rpm for 10 min, and the medium was discarded. Pellets were washed with a S-free buffer of pH 7.0 (Ringer’s), and cells were resuspended in 0.45 ml of 50 mM HEPES buffer, pH 8.0. Suspensions were separated into three equal volume aliquots (0.15 ml) in Eppendorf tubes. Next, 0.15 ml of a 2 mM DBT solution - prepared in the same buffer-were added in each tube, and desulfurization reaction took place under shaking (1200 rpm) for 30 min at 30 °C in a thermostated Eppendorf shaker (Thermo Shaker TS-100, BOECO, Germany). The reaction was terminated with the addition of an equal volume (0.3 ml) of acetonitrile (Labbox Export, Kappa Lab SA, Hellas) and vigorous vortexing. Suspensions were centrifuged (14.000 g; 10 min), and 2-HBP produced was determined in the collected supernatant through HPLC. One of the tubes, where the 0.3 ml acetonitrile was added immediately after DBT addition (t=0), was used as blank. Desulfurization capability was expressed as Units per mg dry cell weight, where 1 Unit corresponds to the release of 1 nmole of 2-HBP per hour. The linearity of the above-described assay with respect to biomass concentration has been verified for up to 2 h reaction time and up to 100 μΜ 2-HBP produced in the sample (Supplementary Figure S4). According to the procedure described above, every technical replicate (N=2) corresponds to the biological mean of 2 - 5 replicate wells.

### HPLC analysis

High-performance liquid chromatography (HPLC) was used to quantify 2-HBP and DBT. The analysis was performed on an Agilent HPLC 1220 Infinity LC System, equipped with a fluorescence detector (FLD). A C18 reversed phase column (Poroshell 120 EC-C18, 4 μm, 4.6×150 mm, Agilent) was used for the separation. Elution profile (at 1.2 ml/min) consisted of 4 min isocratic elution with 60/40 (v/v) acetonitrile/H_2_O, followed by a 15 min linear gradient to 100% acetonitrile. Fluorescence detection was performed with excitation and emission wavelengths of 245 nm and 345 nm, respectively. Quantification was performed using appropriate calibration curves with the corresponding standards (linear range 10 - 1000 ng/ml).

### Extraction of total RNA

*R. qingshengii* IGTS8 wt, *cbsΔ,* and *metBΔ* deletion strains were grown in CDM medium containing DMSO, MgSO_4_, methionine, or cysteine as the sole sulfur source (1 mM), as described in the Materials and Methods section “Growth and desulfurization assays”. Ethanol was used as a carbon source to a final concentration of 0.165 Μ (0.33 M carbon). Cells were harvested in mid-exponential phase and incubated with lysozyme (20 mg/ml) for 2h at 25 °C. Total RNA isolation was performed using NucleoSpin RNA kit (Macherey-Nagel, Lab Supplies Scientific SA, Hellas) according to manufacturer guidelines. RNA samples were treated with DNase I as part of the kit procedure to eliminate any genomic DNA contamination. RNA concentration and purity were determined at 260 and 280 nm using an μDrop Plate with a Multiskan GO Microplate Spectrophotometer (Thermo Fisher Scientific, Waltham, MA USA), while RNA integrity was evaluated by agarose gel electrophoresis.

### First-strand cDNA synthesis

Reverse transcription took place in a 20 μl reaction containing 500 ng total RNA template, 0.5 mM dNTPs mix, 200U SuperScript II Reverse Transcriptase (Invitrogen, Antisel SA, Hellas), 40U RNaseOUT Recombinant Ribonuclease Inhibitor (Invitrogen, Antisel SA, Hellas) and 4 μM random hexamer primers (Takara Bio, Lab Supplies Scientific SA, Hellas). Reverse transcription was performed at 42 °C for 50 min, followed by enzyme inactivation at 70 °C for 15 min. The concentration of cDNA was determined using an μDrop Plate with a Multiskan GO Microplate Spectrophotometer (Thermo Fisher Scientific, Waltham, MA USA).

### Quantitative Real-Time PCR (qPCR)

qPCR assays were performed on the 7500 Real-Time PCR System (Applied Biosystems, Carlsbad, CA) using SYBR Green I dye for the quantification of *dszA, dszB, dszC, cbs,* and *metB* transcript levels. Specific primers were designed based on the published sequences of IGTS8 desulfurization operon (GenBank: U08850.1 for *dszABC*) and IGTS8 chromosome (GenBank: CP029297.1 for *cbs*, *metB, gyrB*) and are listed in Table S1. The gene-specific amplicons generated were 143 bp for *dszA*, 129 bp for *dszB*, 152 bp for *dszC*, 226 bp for *cbs,* 129 bp for *metB* and 158 bp for *gyrB*. The 10 μl reaction mixture included 5 μl Kapa SYBR Fast Universal 2x qPCR master mix (Kapa Biosystems, Lab Supplies Scientific SA, Hellas), 5 ng of cDNA template, and 200 nM of each specific primer. The thermal protocol was initiated at 95 °C for 3 min for polymerase activation, followed by 40 cycles of denaturation at 95 °C for 15 sec, and primer annealing and extension at 60 °C for 1 min. Following amplification, melt curve analyses were carried out to distinguish specific amplicons from non-specific products and/or primer dimers. All qPCR reactions were performed using two technical replications for each tested sample and target, and the average CT of each duplicate was used in quantification analyses, according to the 2^−ΔCCT^ relative quantification (RQ) method. The DNA gyrase subunit B (*gyrB*) gene from strain IGTS8 was used as an internal reference control for normalization purposes. A cDNA sample derived from *R. qingshengii* IGTS8 grown on 1 mM DMSO for 66 h was used as our assay calibrator. All qPCR procedures and results are reported according to the MIQE guidelines. See also Supplementary Table S3.

### Statistical analysis

Specific growth rates (μ_max_) and maximum biomass concentrations (C_max_) were calculated as described in the section “Growth and desulfurization assays”. The results of growth studies (μ_max_ and C_max_ values) were statistically analyzed by performing one sample t-tests for each strain and each medium composition (Confidence interval was set to 95%; N=12-17). For comparison of growth kinetic parameters between different strains or medium compositions, unpaired t-test or one-way ANOVA with Tukey’s Multiple Comparison test was performed depending on the number of compared columns (Confidence interval was set to 95%; N=12-17). Statistical significance is described in the corresponding results sections.

To compare BDS activities (Units/mg DCW) for results shown in Figure 4, one-way ANOVA with Tukey’s Multiple Comparison test was performed between different growth conditions (glucose vs glycerol vs ethanol as sole carbon source). For the statistical analyses of BDS activities in Figures 5, 6, 7, 8 one-way ANOVA with Tukey’s Multiple Comparison test was performed, between the wt and each of the recombinant strains, for the same growth conditions (i.e., same carbon and sulfur sources) at the same time point (20, 45, 65 h). P-values are indicated in the corresponding figures. Our analysis also included comparison of different concentrations of the same sulfur source, supplemented to the same strain. P-values are mentioned in the relevant parts of results. Confidence interval was set to 95% in all cases.

For results shown in Figure 9, one-way ANOVA with Tukey’s Multiple Comparison test was performed, to compare transcription levels of wt and each of the recombinant strains. Biological/Technical replicates: 2/2 (See also Table S3). The software used for statistical analyses was GraphPad Prism 6.01.

## Acknowledgments

We thank Jacob Bobonis (EMBL Heidelberg) for the S17-1 strain. This research project was supported by the Action RESEARCH – CREATE – INNOVATE co-financed by the European Regional Development Fund of the European Union and national resources through the Operational Program “Competitiveness, Entrepreneurship & Innovation” (EPAnEK) - NSRF (2014-2020) (Project code: T1EDK-02074, MIS 5030227).

## Supplementary Legends

**Figure S1. Schematic representation of knockout strain construction.** A constructed vector *pK18mobsacB-5’_TG_-3’_TG_* harboring the upstream (5’ _TG_) and downstream (3’ _TG_) flanking regions of the target gene (TG), is inserted to the bacterial cell of the recipient strain (wt *R. qingshengii* IGTS8) *via* conjugal transfer. In the first step of the gene deletion process, homologous recombination takes place between the transferred DNA (vector) and the chromosomal DNA of the recipient strain, resulting in either 5’ (upstream) or 3’ (downstream) plasmid incorporation. The generated transconjugants are kanamycin resistant (Kan^R^) and sucrose sensitive (Suc^S^), allowing for kanamycin resistance selection. As an additional control, transconjugants should exhibit no growth in the presence of 10% sucrose. The validated upstream or downstream merodiploids, are grown in the absence of selective pressure (without antibiotic addition) to induce the second homologous recombination event. Depending on the flanking regions involved (5’ or 3’), the result is either a knockout of the target gene *tgΔ* or a reversion to the wt, while the rest of the integrated vector is excised from the genome. The generated strains are sucrose resistant, thus any incomplete excision events are removed in the final step of the process, with selection in the presence of 10% sucrose. Additionally, kanamycin sensitivity is restored at this point. Single colonies are isolated, and knockouts are confirmed with PCR using 5F-check and 3R-check primer pair. The PCR products are further verified with DNA sequencing.

**Figure S2. Growth and 2-HBP production in the presence of DBT as sole sulfur source.** Growth (Biomass; g/L) and 2-HBP concentration in the culture medium of wt, *cbsΔ*, and *metBΔ* isogenic strains in the presence of 0.1 mM ***DBT*** as sole sulfur source. Ethanol (0.165 M) was supplemented as the sole carbon source. For measurements of produced 2-HBP (μΜ) at 0, 24, 48, and 72 h, an aliquote of whole culture was harvested in eppendorf tubes and an equal volume of acetonitrile was added. Suspensions were vortexed vigorously, centrifuged (14.000 g; 10 min), and 2-HBP concentration was determined in the collected supernatant through HPLC. Calculated μ_max_ and C_max_ values are included in the upper left inset (n=3).

**Figure S3. Effect of cysteine supplementation at a high concentration.** Growth curves (Biomass; g/L) and desulfurization activities (Units 2-HBP/mg DCW) of wt, *cbsΔ*, and *metBΔ* isogenic strains in the presence of 10 mM ***cysteine*** as sole sulfur source. Ethanol (0.165 M) was supplemented as the sole carbon source in the culture medium. Calculated μ_max_ and C_max_ values are included in the upper left inset.

**Figure S4. Linearity of BDS assay with respect to biomass concentration.** Graph showing the linear relationship between the concentration of detected 2-HBP (μM) and the concentration of biomass (g/L) used in the BDS assay. The line corresponds to linear regression for the plotted points. Assay time 1 h.

**Table S1.** Oligonucleotides used in this study.

**Table S2.** Growth kinetic parameters (μ_max_ and C^x^_max_) obtained by fitting of biomass concentration vs. time experimental values to the logistic equation.

**Table S3.** Datasheet for MIQE guidelines.

**Supplementary text.** DNA sequences of *cbs-metB* genetic loci for wild-type and recombinant strains.

